# Cancer cachexia overrides sex-specific traits in Drosophila

**DOI:** 10.64898/2026.10.09.757864

**Authors:** Alexia Rivero, Margaux Techer, Nourhene Ammar, Hadi Boukhatmi, François Payre, Jennifer Zanet, Cédric Polesello

## Abstract

Cancer cachexia, a metabolic syndrome characterized by pathologic weight loss, dramatically limits therapeutic options and reduces survival. While the mechanisms by which tumors induce wasting of distant organs (particularly muscle and adipose tissues) remain incompletely understood, the influence of biological sex on cachexia progression is even less clear. Here we investigated phenotypic and molecular manifestations of cachexia in both sexes using a model of inducible intestinal tumors in *Drosophila*. We show that tumor progression leads to the development of severe cachexia both in males and in females manifested by organ atrophies and an inflammatory state. Nevertheless, both phenotypical and molecular cachexic alterations exhibit striking differences between sexes. Transcriptomic analysis show that many genes differentially expressed at high levels in males under normal conditions become downregulated during cachexia, while their expression is upregulated in cachexic females, and reciprocally. Together, sex-specific deregulation in gene expression and metabolic profiles ultimately converge towards a molecular cachexic state that erase or diminish many sex-specific differences otherwise characteristic of healthy males and females. These findings highlight the interplay between oncogenic stress and biological sex in shaping cachexia phenotypes, shedding light on how sex influences the development of cancer cachexia.

## Introduction

Sexual dimorphism has a major effect on cancer incidence and patient survival. Non-reproductive cancers are more prevalent in men, who also have a mortality rate nearly twice that of women (Clocchiatti *et al*, 2016). This disparity persists even after accounting for differences in environmental exposures (*e.g.*, smoking, alcohol consumption, …) and risk behaviors (Dorak & Karpuzoglu, 2012) highlighting the intrinsic role of biological sex in cancer susceptibility (Dorak & Karpuzoglu, 2012). In addition to tumor onset and progression, biological sex also influences the systemic complications resulting from cancer.

Cachexia, a multifactorial inflammatory syndrome, accompanies approximately 50% of cancer cases and causes dramatic weight loss, including skeletal muscle atrophy and adipose tissue depletion (Ferrer *et al*, 2023). Its progresses through distinct stages: pre-cachexia, cachexia, and terminal cachexia, correlating with the extent of weight loss (Watanabe & Oshima, 2023). Unfortunately, cachexia is often diagnosed at late stages, compromising the efficacy of anti-cancer surgery and chemotherapy and thus reducing the chances of remission. Notably, the incidence and severity of cachexia vary according to both cancer type and patient sex. Males are more likely to develop cachexia, often with greater intensity (Stephens *et al*, 2012), as seen from a higher proportion of muscle loss (Wallengren *et al*, 2015; Norman *et al*, 2012) and a reduced quality of life (Kilgour *et al*, 2010). Several hypotheses have been proposed to explain these sex-based disparities in cachexia. First, the ratio of slow (type 1) to fast (type 2) muscle fibers differs between sexes, men have more fast fibers and are more susceptible to cachexia (Acharyya *et al*, 2005). Second, female hormones, particularly estrogens, may exert protective effects on skeletal muscle integrity (Kitajima & Ono, 2016). Finally, sex differences in the inflammatory response, particularly involving Interleukin 6 (IL6) that is a key mediator of cachexia, may contribute to these disparities. For instance, in mice, muscle cells in females express fewer IL-6 receptors and, consistently, appear less sensitive to circulating IL-6 levels (Hetzler *et al*, 2015).

Given the challenges of early diagnostic of cachexia in humans, animal models have been developed to elucidate the mechanisms underlying this syndrome (Ferrer *et al*, 2023). Among these, the inducible manipulation of intestinal stem cells (ISCs) in flies has provide a valuable paradigm to study cancer cachexia. Expression of an activated form of the Yorkie oncogene (Yki^act^(Oh & Irvine, 2009)) in ISCs induces intestinal hyperplasia, followed by progressive degeneration of muscle and adipose (fat body) tissues, as well as kidney dysfunction leading to abdominal bloating, thereby mimicking aspects of human cachexia (Ferrer *et al*, 2023). As also observed in humans, cachexia progression in flies is accompanied by metabolic disorders, including defects in glucogenesis, tricarboxylic acid (TCA) cycle, as well as synthesis of Trehalose, the main sugar present in the fly blood stream (Liu *et al*, 2025). Induction of the cachectic state involves four factors that are released by tumorous intestines into the circulatory system, called hemolymph in flies. They comprise Impl2 (the functional equivalent of IGFBP7) that binds insulin-like peptides (Ilps) and inhibits insulin signaling in peripheral tissues (Kwon *et al*, 2015). Second, PVF1, a PDGF-VEGF-related factor, activates the MEK signaling pathway, leading to increased muscle catabolism and impaired lipid storage (Song *et al*, 2019). Third, Upd3, the ortholog of the inflammatory marker IL6, activates the JAK/STAT pathway, promoting both the proliferation of ISCs and the transcription of Impl2 in muscles and fat body (Ding et al., 2021). The secretion of Impl2 by target organs therefore reinforces inhibition of the insulin pathway and increases organ waste (Ding *et al*, 2021). Finally, the anti-diuretic hormone Ion transport peptide (ITP) alters osmotic regulation in Malpighi tubules (the functional equivalent of mammalian kidneys). ITP acts thought binding to its receptor TkR99D present on the surface of renal stellate cells, thereby disrupting the osmotic equilibrium that results in a prominent abdominal bloating characteristic of cachectic flies (Xu *et al*, 2023).

However, *Drosophila* cancer cachexia has mainly been studied in females, since mated female flies are well-known to display increased intestinal cell renewal rate that facilitates tumor induction (Hudry *et al*, 2016a). In this study, we examined the influence of sex on cachexia and its phenotypic manifestations following the expression of Yki^act^ in ISCs. We found that Yki^act^ induces tumorigenic over-proliferation of ISCs in males and females, ultimately leading to cachexia in both sexes. Despite initial sexual dimorphisms, our transcriptomic analyses reveal that males and females yet converge toward a similar molecular signature characteristic of the cachexic state, underscoring the complex mechanisms underlying cancer cachexia.

## Results

### Male intestines develop tumors under Yki gain of function in intestinal stem cells

Intestinal homeostasis is maintained by intestinal stem cells (ISCs), which are capable of renewing all cell types of the intestine, including differentiated enterocytes (ECs) and enteroendocrine cells (EEs) (Ohlstein & Spradling, 2006; Micchelli & Perrimon, 2006) (Fig 1A). This differentiation occurs via progenitor stages: enteroblasts (EBs) and pre-EE cells for ECs and EEs, respectively (Zeng & Hou, 2015). The *Drosophila* intestine shows several disparities between males and females, including spatial layout in the body (Blackie *et al*, 2024), metabolism (Hudry *et al*, 2019), and proliferative activity of stem cells (Hudry *et al*, 2016b). Once mated, the metabolism of females changes radically to support oogenesis, which requires a strong increase in nutrient intake and thus represents a challenge to maintain the gut integrity (Cognigni *et al*, 2011; Reiff *et al*, 2015; O’Brien *et al*, 2011). Hence, mated females display higher metabolism and ISC renewal, when compared to virgins (Lee *et al*, 2013; Carvalho *et al*, 2006). Turnover of male intestinal tissue is also far slower and male ISCs are generally thought to be quiescent. Accordingly, male ICS have been reported to be less prone to intestinal tumors, for example upon deregulation of the Notch signaling pathway (Hudry *et al*, 2016b). Indeed, most studies on intestinal tumors and cachexia have hitherto been conducted only in female flies.

**Figure 1:**
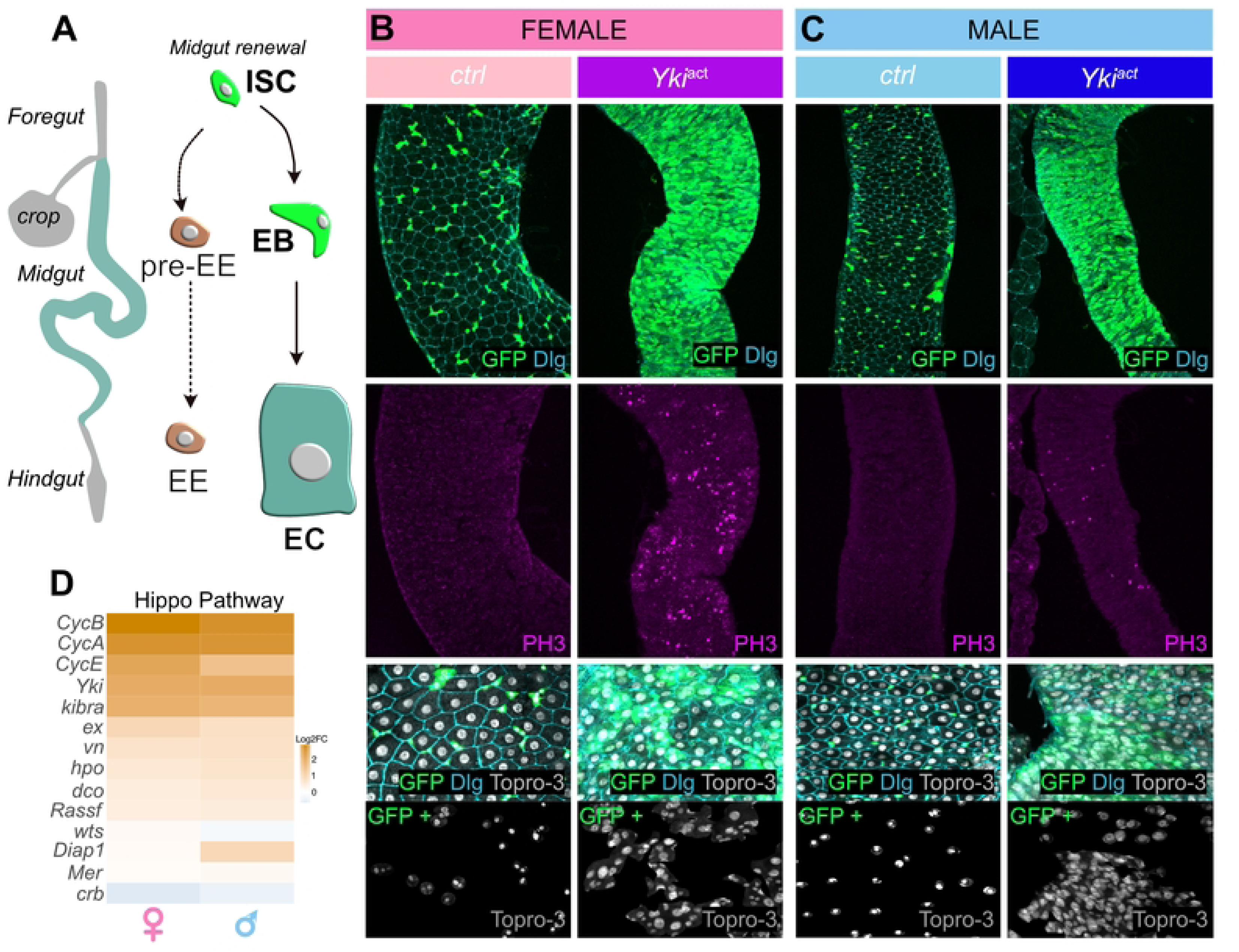
Yki^act^ promotes tumor growth in the intestine of both males and females. **(A).** Schematic representation of the adult *Drosophila* intestine (left) and stem cell lineage (right). In the midgut, intestinal stem cells (ISCs) are capable of regenerating both enteroendocrine cells (EEs) and enterocytes (ECs) via progenitor stages, called pre-EEs and enteroblasts (EBs), respectively. **(B-C).** Images of the midgut region of the female **(B)** or the male **(C)** intestine. GFP alone (controls) or in combination with Yki^act^ was expressed with *esg^ts^* (*w; esgGAL4, UAS-GFP; tub-GAL80ts*). The upper panel shows the distribution of ISCs, as revealed by GFP. Mitotic figures are identified in the middle panels by staining for the phosphorylated form of Histone 3 (pH3, purple). The bottom panels illustrate the heterogeneity of nuclei of ISC/EB cells in Yki^act^ conditions. Membranes were stained using Disc large (Dlg, cyan) and nuclei by Topro-3 (grey). **(D).** Heatmaps representing expression dynamics (log2(FC)) of known targets of the Hippo pathway in female and male tumorous intestines.

Given the influence of biological sex in cancer cachexia in humans, we looked for a suitable model system in flies. During the course of our studies, we found that, unlike Notch manipulation, an active form of Yki drives intestinal tumors in both sexes, with phenotypes indicative of cachexia in males and females. We used the *UAS/escargot-Gal4/Gal80* thermosensitive system (referred to as *esg^ts^* (Jiang *et al*, 2009)) to express Yki^act^ (Oh & Irvine, 2009) and GFP in ISCs. Consistent with previous findings (Kwon *et al*, 2015; Liu *et al*, 2025), females intestines displayed massive accumulation of GFP-positive cells due to ISC over-proliferation, as evidenced by the increase in PH3-positive cells (Figure 1B). ISC over-proliferation was detectable as early as five days post-induction, and persisted at least to fifteen days (Fig Supp1). We found that males showed a similar pattern (Fig 1C and Fig Supp1), with tumor growth (S1 Fig) and increased cell division (Fig 1B).

In females, it has been reported that Yki^act^ expression in ISC/EB cells induces abnormal heterogeneity in the enterocyte lineage, which results in an increase and larger variability in nuclear size (Pranoto et al., 2023; and Fig 1B). The same phenotypic manifestation of Yki^act^ in ISC/EB cells was also observed in males (Fig 1C). These results show that Yki^act^ expression in males or females ISCs induces intestinal tumors, which display similar cellular features in both sexes.

To analyze the transcriptional response following expression of Yki^act^, we performed RNA-sequencing (RNA-seq) on whole intestines from both sexes, ten days after induction, when cachexia is well developed in females (Kwon *et al*, 2015). Examination of target genes confirmed the activation of Hippo signaling in male and female intestines (Fig 1D). Principal component analysis (PCA) well separated global gene expression profiles between control and Yki^act^ conditions, for both sexes (Fig 2A). Analysis of gene expression showed that ISC/EB markers were overrepresented in both sexes under Yki^act^ conditions, whereas markers for mature ECs were underrepresented (Fig 2C). The data then support that tumoral condition has a larger effect than the sex on intestinal gene expression.

**Figure 2:**
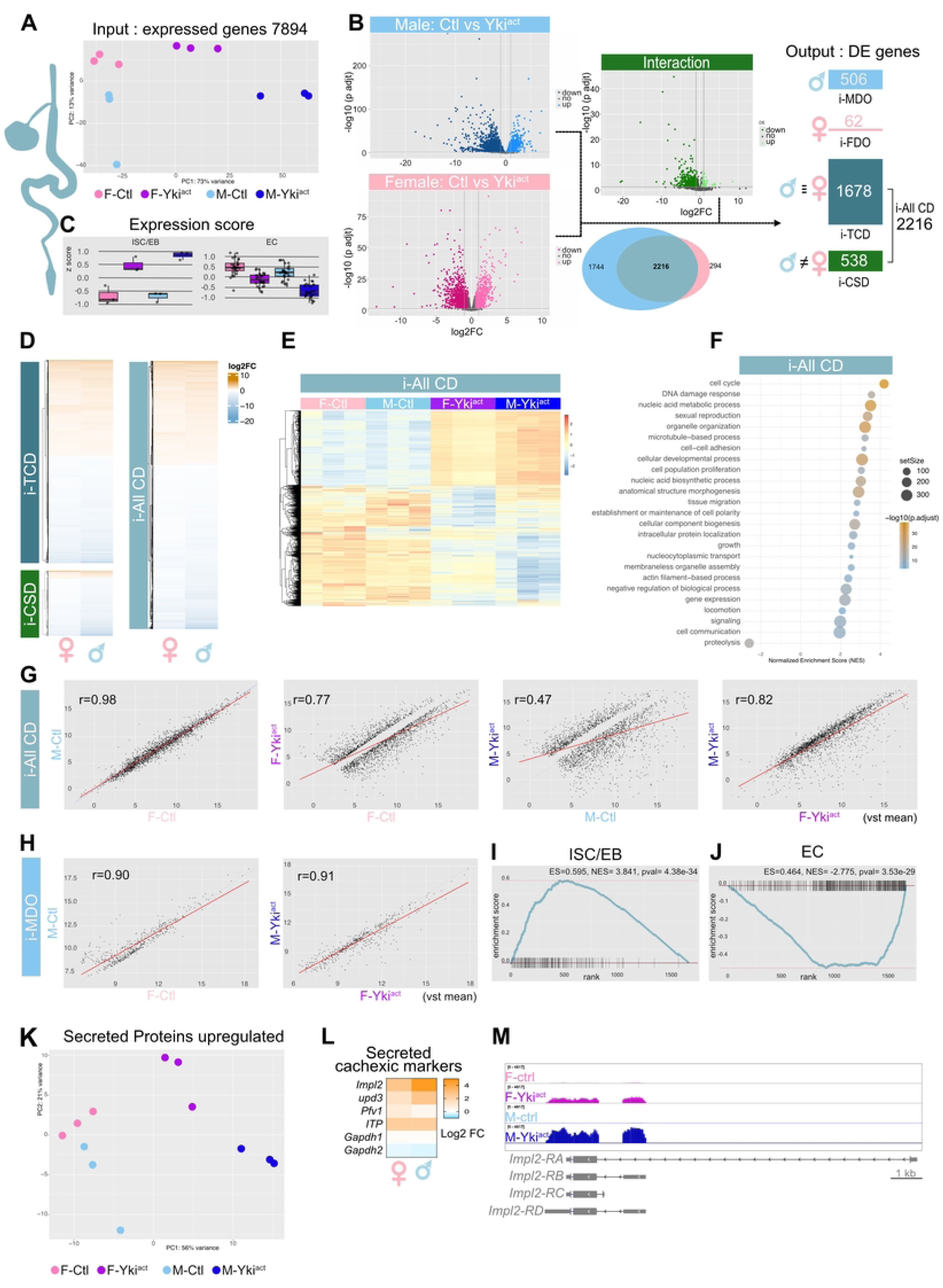
Intestinal tumors in females and males exhibit similar gene expression dysregulation. **(A).** Principal component analysis (PCA) of all genes expressed in the intestine based on RNA seq data, after ten days of transgene induction. **(B).** Volcano plots (left) show differentially expressed genes (DEGs) upon Yki^act^ expression in adult ISCs at ten days, in females (pink) and males (blue). The Venn diagram shows intersection of the two volcano plots. 2216 genes are commonly deregulated (i-AllCD) in males and females following Yki^act^ expression. The volcano plot shown in green, called “interaction,” identifies DEGs taking into account both sex-related effects and Yki^act^ effects. The combination of these two methods identified four groups of differentially expressed genes: i-MDO (intestinal Male Deregulated Only), i-FDO (intestinal Female Deregulated Only), i-TCD (intestinal Truly Commonly Deregulated), and i-CSD (intestinal Common but Sex Dependent). **(C).** Expression scores for cell-type-specific markers of ISC/EBs and ECs, in control and Yki^act^ conditions, for females and males, using the same color code than in **(A)**. **(D).** Heatmap showing the behavior (log2 FC) of the groups i-TCD, i-CSD and i-AllCD. **(E).** Heatmap of the expression of 2216 genes included in the group i-AllCD according to sex and oncogenic conditions (control *vs* Yki^act^). **(F).** Dot plot showing the results of Gene Set Enrichment Analysis (GSEA) that illustrate the biological processes (GO terms) associated with i-AllCD genes. GeneRatio represents the number of input genes mapped to a given GO term divided by the total number of input genes. Statistical significance was assessed by multiple comparisons using the Benjamini Hochberg method. KEGG, Kyoto Encyclopedia of Genes and Genomes. **(G).** Scatter plots illustrating the correlation between the expression levels of DEGs in the i-AllCD group, comparing sex by condition and condition by sex. **(H).** Scatter plots testing the correlation between sex by condition for the i-MDO group. **(I-J).** GSEA of the ISC/EB (**I**) or EC signatures (**J**), performed on group i-TCD genes. **(K).** PCA of all genes encoding putatively secreted proteins (*i.e.*, containing a signal peptide) for each condition and sex. **(L).** Heatmap showing expression dynamics (log2 FC) of genes encoding secreted molecules identified as impacting the development of cachexia. Gapdh1 and Gapdh2 were used as controls. **(M).** Genome browser screenshot of the *Impl2* locus (control females in pink, Yki^act^ females in purple, control males in light-blue, Yki^act^ males in dark blue).

We next sought to identify differentially expressed genes (DEGs) in Yki^act^-expressing intestines, separately for each sex (Fig 2B). 3,960 DEGs were identified in males and 2,510 in females, with an overlap of 2,216 genes deregulated in both males and females (Fig 2B, Venn diagram). We conducted a complementary analysis, which took into account the interaction between sex, tumoral conditions and a significant effect of Yki^act^ (Fig 2B, and see Methods). By intersecting the two types of analyses, we identified four groups of genes that are dependent on Yki^act^ expression in ISCs (Fig 2B, output DE genes). Genes in groups i-TCD (intestine Truly & Commonly Dysregulated, 1678 genes) and i-CSD (intestine Common but Sex Dependent, 538 genes) are deregulated in both sexes, whilst levels of deregulation differ between sexes in the latter group. i-MDO (Intestine Male Dysregulated Only, 506 genes) corresponds to DEGs in Yki^act^ intestines that are specific to males; i-FDO (intestine Female Deregulated Only, 62 genes) to DEGs specific to females. Further analysis of the i-TCD and i-CSD groups revealed that gene deregulation follows a similar pattern in both sexes, although i-CSD genes show a more pronounced deregulation in males (Fig. 2D). We grouped these genes into a single cluster, “i-AllCD” (intestine All Commonly Dysregulated genes), whose behavior is illustrated in Fig 2E, D. Approximately 40% of these genes were overexpressed under Yki^act^ conditions. Gene set enrichment analysis (GSEA) linked a significant proportion of them to cell cycle progression (Fig 2F), which is consistent with the observed excessive proliferation (Fig 1B, C). Markers of ISC/EB are overrepresented, while those for ECs are underrepresented (Fig. 2I, J). The i-AllCD set of genes, which accounts for approximately 80% of DEGs for the four groups, exhibits similar basal levels in males and females (correlation coefficient, r =0.98, Fig 2G). This confirms that the intestines of both sexes are very similar. Yki^act^ expression in ISCs influences the expression of these genes, particularly in males (r=0.47), but their expression levels remain closely correlated under Yki^act^ conditions (*r* ≈0.8, Fig 2G). This suggests that Yki^ACT^ influences gene expression of i-AllCD set in a similar manner between the two sexes, increasing ISCs/EBs and reducing differentiated ECs (Fig 2I, J). Genes upregulated only in males show similar correlation coefficients in control and Yki^ACT^ conditions (*r* ≈0.9), further supporting the similarity between male and female intestinal responses (Fig 2H). Since the onset of cachexia is induced by secreted molecules produced by tumorous intestines (Liu *et al*, 2022), we studied the expression of DEGs encoding proteins with a signal peptide (identified in FlyBase (S1 Table). As observed for the whole set of genes (Fig 2A), genes encoding secreted molecules easily separate control from oncogenic intestines, for both sexes (Fig 2 K). This behavior is well exemplified by the four secreted-molecules (*Impl2*, *Upd3*, *Pvf1*, and *ITP)* previously shown to contribute to cachexia development in females (Kwon *et al*, 2015; Ding *et al*, 2021; Song *et al*, 2019; Xu *et al*, 2023). After ten days of oncogenic stress, the four genes encoding these diffusible proteins were strongly upregulated in the intestines of females and males (Fig 2L, M). Thus, in addition to exhibiting phenotypic behavior similar to that of females, the tumor-bearing intestines of males also produce molecules involved in the onset of cachexia.

Overall, the intestinal response to oncogenic Yki^act^ appears strikingly similar between sexes; including cell phenotypic changes, excessive stem cell proliferation, as well as transcriptional dysregulation leading to the production of signaling molecules.

### Signs of cachexia appear in males with intestinal tumors

Having shown that Yki^act^ induces intestinal tumors in both sexes, we sought to determine whether this oncogenic stress could also trigger cachexia in males, as documented in females (Kwon *et al*, 2015). We first investigated phenotypic manifestations of cachexia, including adult survival, renal function and muscle activity. Under our standard growth conditions, healthy male and female flies exhibited a comparable lifespan, surviving up to forty-five days (Fig 3A). Flies with intestinal tumors showed reduced lifespan, with males and females succumbing approximately twenty days earlier than their wild-type siblings (Fig 3A). Disruption of the renal function is a hallmark of cachexia in flies, leading to prominent abdominal bloating (Kwon *et al*, 2015). Females displaying abdominal bloating were observed as early as five days after Yki^act^ induction, reaching sixty percent of the population by day ten (Fig 3B). Males also developed abdominal bloating, although with a delay of a few days compared to females. The first signs of bloating appearing by day ten (Fig 3B) and reached about fifty percent by day nineteen. We further observed other evidence of renal dysfunction in both sexes, even if these defects were more pronounced in females. These included a decrease in the number of renal stem cells, as well as the formation of kidney stones containing calcium oxalate, calcium phosphate, and xanthine crystals (Reynolds *et al*, 2021), these abnormalities being more pronounced in females (Fig 3D). Finally, we assessed muscle activity using a climbing test, measuring the fly ability to scale back to the top of their vial after tapping. This indirect test has been previously employed to characterize tumor-induced cachexia in females (Kwon *et al*, 2015). Consistently, we observed significant climbing defects in females, with approx. 70% failure rate at fifteen days (Fig 3C). Although males might exhibit a similar trend, the increase in climbing failure was not statistically significant (Fig 3C). Of note, under our experimental conditions of cachexia, we failed to detect obvious abnormalities in the architecture of indirect flight muscles, regardless of sex (Fig 3E).

**Figure 3:**
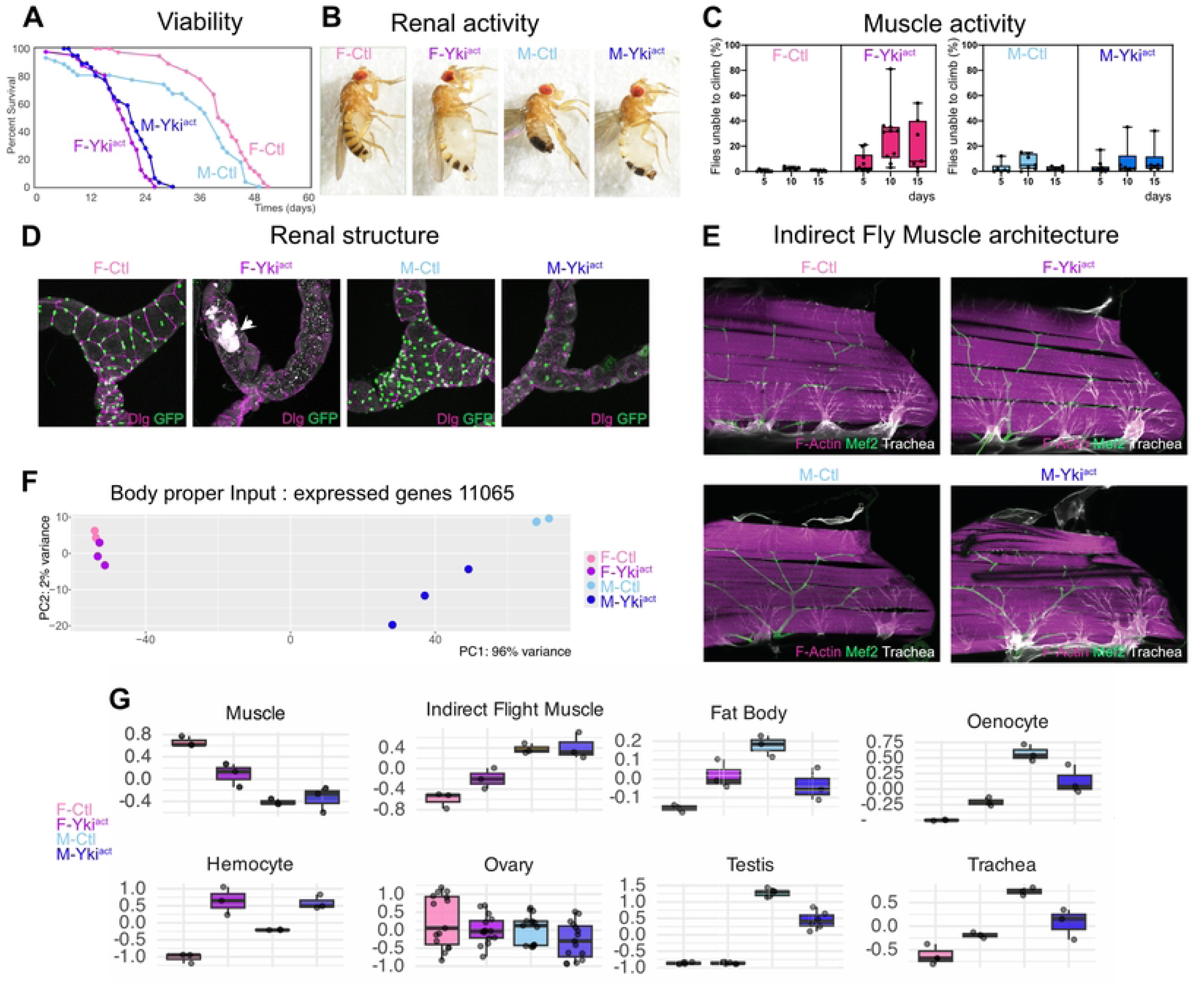
Effects of intestinal tumors on the body proper. **(A).** Survival curve of male and female flies under control and Yki^act^ conditions. **(B)**. Pictures of adult flies, at ten days, representing males and females in control and Yki^act^ contexts. Note the bloating abdomen seen in both sexes after ten days of Yki^act^ expression. **(C)**. Time course of climbing tests in males and females expressing or not Yki^act^ in adult ISCs. **(D)**. Images of adult Malpighian tubules, which are responsible for renal function in flies. Renal and nephric stem cells (RNSCs) are visualized by GFP expression (green) expressed by *esg^ts^*. Septate junctions were stained with anti-Disc Large (Dlg) antibody (magenta). Kidney stones are visible by auto-fluorescence (grey arrowhead). Tubules were analyzed after ten days of transgene induction. **(E)**. Longitudinal optical section of indirect flight muscles of adult females (top panels) or males (bottom panels) in control (left) and Yki^act^ backgrounds, ten days after transgene induction. F-actin is shown in pink, Mef2 that highlights muscle nuclei in green, and tracheal cells are in gray. **(F)**. PCA of all genes expressed in the body-proper (*i.e.,* all tissues after removal of the head and intestine) after ten days of induction (GFP alone or in combination with Yki^act^). **(G).** Boxplots illustrating the expression dynamics of gene sets characteristic of the indicated tissues. Quantification was performed using RNA-Seq data from adult body samples in males and females, from control and Yki^act^ conditions.

Taken together, these results indicate that males with Yki^act^-induced intestinal tumors exhibit multiple signs of cachexia, including reduced lifespan and renal dysfunction. However, the severity of these phenotypic manifestations appears less pronounced than in females, particularly with regards to muscle activity. This suggests that although cachexia occurs in both sexes, the frequency and severity of cachexia in flies is influenced by the sex of tumor bearing animals.

### Assessment of the molecular signature of cachexia in target tissues

To investigate the transcriptional consequences of tumor-induced cachexia in target tissues, we performed RNA-seq on the remaining body tissues after dissecting off the head and intestine, hereinafter referred to as “body proper” samples. Body-proper RNA-seq was run for controls and animal bearing Yki-induced intestinal tumors, in male and female individuals. As expected, since non intestinal tissues display obvious morphological differences between males and females, PCA detected a marked distinction between male and female body-proper transcriptomic profiles, both in control and tumorous individuals (Fig 3F, compare to Fig 2A). This global view of gene deregulation in target organs of animals with intestinal tumors yet suggested that transcriptional changes were more pronounced in males than in females (Fig 3F). To further investigate this possibility, we analyzed gene expression of organ-specific markers, as defined by single-cell transcriptomic data (Li *et al*, 2022), across different target tissues, and for each sex (Fig 3G). In females, we confirmed that Yki^act^ intestinal tumors triggered transcriptional changes in primary target organs of cachexia, *i.e.,* fat body, muscles (including indirect flight muscles), and ovaries (Known et al 2015). In addition, the results suggested that oenocytes, hemocytes, and tracheal cells may also be affected by cachexia (Fig 3G). In males, we similarly observed gene deregulation in fat body, hemocytes, tracheal cells, oenocytes, as well as in testes. However, the deregulation of muscle markers was barely detectable in males (Fig 3G), which may explain why Yki^act^ males performed better than females in climbing assays used to assess muscle activity.

Taken together, these results support that Yki^act^ intestinal tumors elicit phenotypic and molecular manifestations of cachexia, in both males and females, with sex-specific differences in the target organs, including magnitude and/or timing of these defects.

### Inflammation as a common feature of male and female cachexia

To identify DEGs in target tissues under intestinal oncogenic stress, we followed the same analytical pipeline as previously described for the intestines (Fig 4A). Comparison of control and Yki^act^ conditions in either males or females indicated that levels of gene deregulation are globally more pronounced in males (4,658 DEGs in males compared to 2,642 DEGs in females, Fig 4A). After intersecting with the results of interaction analyses, we retrieved four distinct groups of DEGs, analogous to those characterized in the intestine (Fig 4A). The body-TCD group (514 genes) and body-CSD group (386 genes) are deregulated in both sexes, although the magnitude of deregulation differs between males and females for the body-CSD group (Fig 4B). The body-MDO group (1,918 genes) comprises male-specific DEGs, while the body-FDO group (1,142 genes) represents female-specific DEGs (Fig 4A). In body-proper samples, the proportion of genes co-deregulated in both sexes was only 23% (compared to 80% in the intestine), supporting the notion that target tissues affected by cachexia display larger sex-specific differences than intestinal tumors.

**Figure 4:**
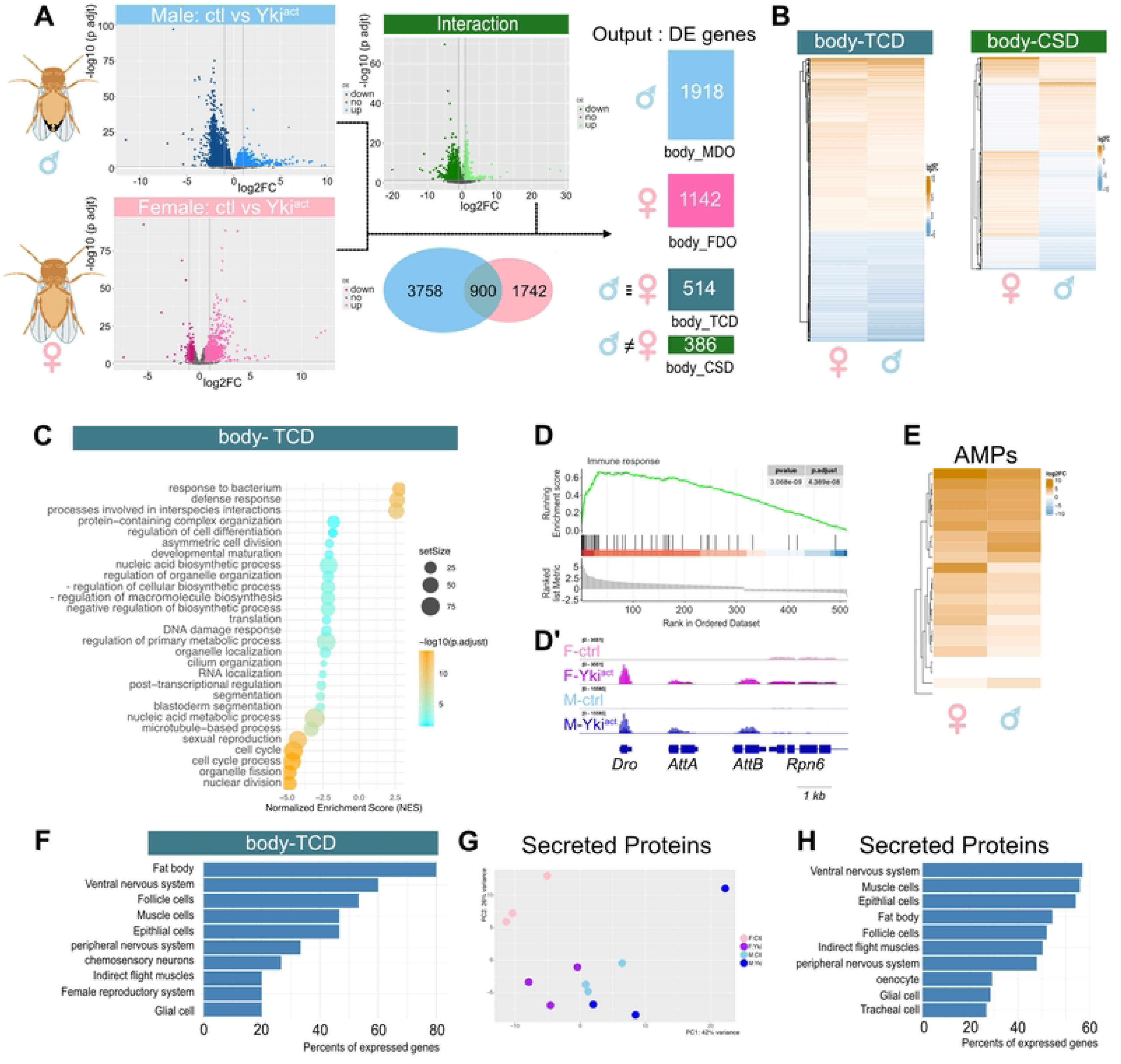
Oncogenic stress triggers an inflammatory response in both sexes. **(A).** Volcano plots show DEGs in body proper samples upon Yki^act^ expression in adult ISCs for ten days; females are in pink (bottom), and males in blue (top). The Venn diagram shows the intersection of volcano plots from males and females. The green volcano plot (interaction) represents DEGs identified by combining the effects of sex and Yki^act^. These analyses allow the identification of four group of DEGs. Groups body-TCD and body-CSD represent genes deregulated in both males and females following Yki^act^ expression. For genes in group body-CSD, levels of their regulation are sex-specific. The group body-FDO represents female-specific DEGs, and body-MDO male-specific ones. **(B).** Heatmaps showing the expression (log2 FC) of genes for body-TCD and body-CSD groups. **(C).** Dot plots of GSEA analysis performed on the body-TCD gene set. **(D).** GSEA on immune response genes (GO:0006955), performed on all DEGs. **(D’).** Genome browser snapshot illustrating the expression of AMPs encoding genes (*Dro*, *AttA* and *AttB*), in the two sexes and genetic contexts. **(E).** Heatmap showing expression dynamics of AMP-coding genes, in body proper samples. **(F).** Percentage of expression of various tissue-specific markers, as defined by (Li *et al*, 2022), in the body-TCD group.**(G)**. PCA showing the behavior of genes encoding putatively secreted proteins (listed in Flybase as containing a signal peptide motif), according to sex and Yki^act^ conditions. **(H).** Percentage of markers’ expression for genes encoding putatively secreted proteins.

We focused on composition and behavior of the body-TCD group, which displayed comparable expression dynamics in both sexes (Fig 4B). We found that the set of body-TCD genes are mostly expressed in muscles, fat body, and epithelial cells (Fig 4F), consistent with the fact that muscle and adipose tissues are being deeply affected by cachexia. By exploring gene ontology data, we noticed that the top three categories were related to the defense response (Fig 4C). This suggests that genes involved in the immune and inflammatory response are the most strongly deregulated in both sexes. Our results also revealed the up-regulation of marker genes characteristic of hemocytes (Fig 3G). Hemocytes represent the main population of immune cells in *Drosophila*, suggesting that an increase in the abundance of hemocytes could contribute to exacerbate the inflammatory state curing cachexia progression. In addition, GSEA analysis performed on all DEGs of body-proper samples showed a clear enrichment of immunity-related genes in cachectic conditions (Fig4 D, D’). A study has recently identified a number of antimicrobial peptides (AMPs) circulating in the body of cachectic flies (Singh *et al*, 2025). Accordingly, we observed overexpression of numerous genes encoding AMPs under cachectic conditions in both sexes (Fig 4E), which could be induced by the fat body (Fig 4F), one of the main sources of AMPs (Lemaitre & Hoffmann, 2007), whereas we did not observe AMP expression in the intestine. In contrast, PCA didn’t detect a particular trend for all candidate genes to encode secreted molecules, which with are expressed from a wide variety of tissues (Fig 4G, H).

Taken together, these data indicate that a common aspect of cachexia in both males and females is the upregulation of genes involved in immune defense, and inflammatory responses to tumors.

### Oncogenic stress induces a comparable cachectic state in males and females

During our analysis of global gene expression, we noticed that body-proper samples showed greater similarity between males and females under cachectic than control conditions. For example, the correlation coefficient for comparing the two sexes was smaller under control (r=0.40) than under cachexia (r=0.75) conditions (S2 Fig). These findings suggest that cachectic males and females display closer gene expression profiles than their healthy counterparts.

To explore this hypothesis, we analyzed the correlation in gene expression between males and females for each of the four gene groups, comparing control and cachectic states. For the body-CSD group (Fig 5A-D), the correlation between males and females is very weak under control conditions (r = 0.26, Fig 5B-C), and gene deregulation strongly varies between sexes, both in intensity and in direction (up *vs* down) of changes (Fig 5A). Some sex-specific deregulations even exhibit antagonism behavior, being overexpressed in one sex and downregulated in the other, or vice versa (Figs 4B, 5A). Interestingly, the correlation coefficient of their expression increases to 0.75 under cachectic conditions (Fig 5C’), suggesting that gene expression profiles progressively converge to a common state between males and females, as the intestinal tumor progresses. This is well exemplified by *geko*, which, like many genes in this group, exhibits opposite deregulation between males and females, ultimately reaching more comparable expression levels under cachectic conditions than in controls (Fig 5D). This general trend was also detected by PCA and correlation matrix analyses (Fig 5B, I). Surprisingly, a similar pattern was also observed for genes in the body-MDO (Fig 5E-G’, J) and body-FDO groups (Figs 5K, S2 Fig), which are differentially regulated according to sex. For the body-TCD group, the correlation was more similar (r=0.63 vs 0.77) for controls and cachectic animals (Figs 5H, S2 Fig), as also seen in PCA (S2 Fig). Hence, as expected for this group of genes that includes AMPs, males and females display comparable responses to intestinal tumors.

**Figure 5:**
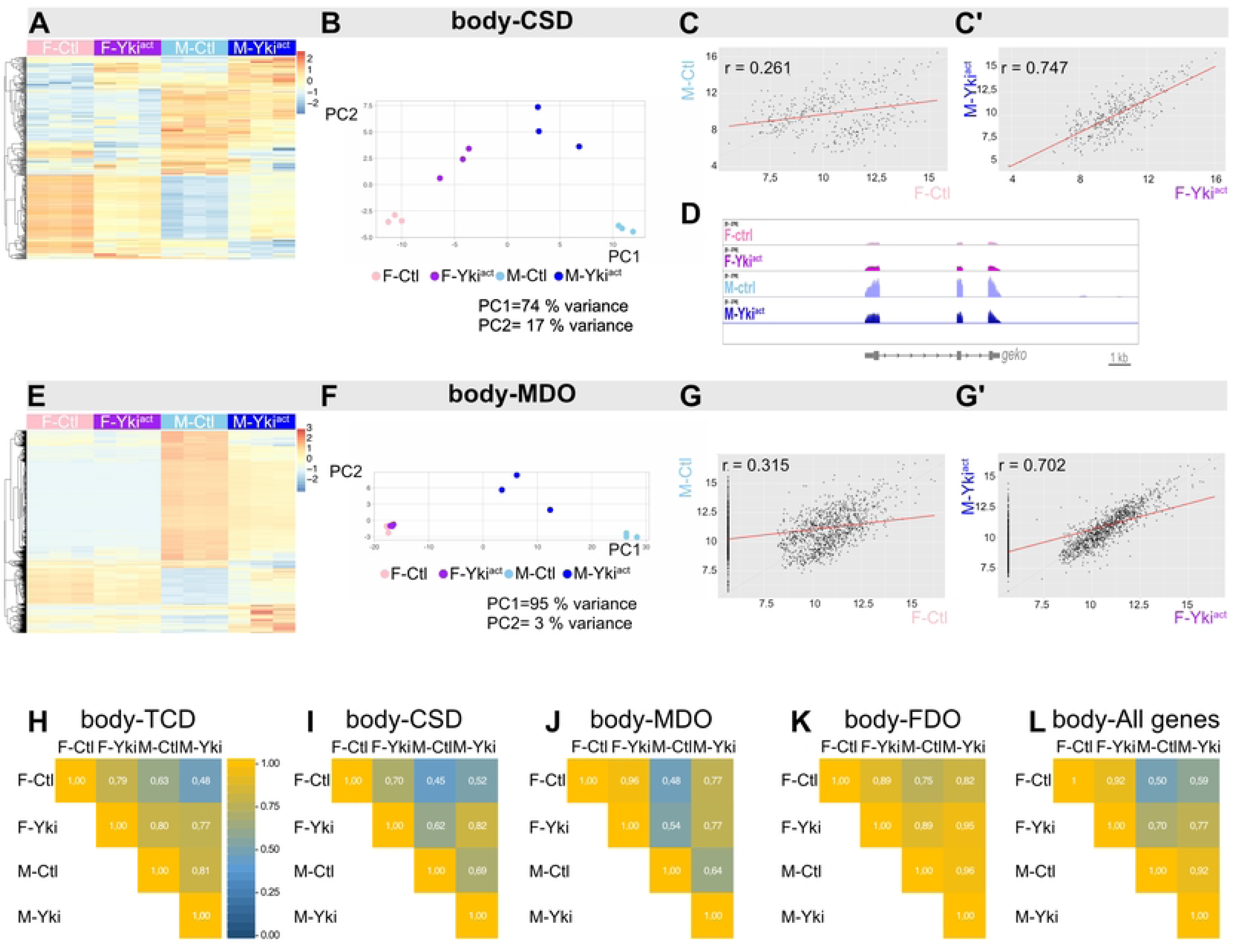
Oncogenic stress causes target organs of males and females to converge towards a similar cachectic state. **(A, E)**. Heatmaps showing overall expression dynamics of body-CSD **(A)**, and body-MDO **(E)** groups of genes, in control and Yki^act^ conditions for both sexes. **(B, F)**. PCA of genes included in the indicated groups. **(C, G)**. Scatter plots showing the correlation between expression levels of body-CSD **(C, C’)** and body-MDO **(G, G’)** groups of genes, in control and Yki^act^ conditions for both sexes. **(D).** Screenshot of genomic browser showing the *geko* locus, illustrating a typical behavior for body-CSD genes. **(H-L).** Gene expression correlation matrixes for each group of body-proper samples, as indicated at the top of each chart.

Taken together, these results support the conclusion that the expression levels of differentially expressed genes converge in cachectic flies of the two sexes. During cancer cachexia, sex-specific molecular deregulations ultimately lead to a similar pattern of gene expression that irons out transcriptional sex-dimorphism.

### Oncogenic stress leads males and females into a similar metabolic state

As observed in mammals, the onset of cachexia in female flies is associated with significant alterations in metabolic processes. For instance, abnormalities in glycolysis, Krebs cycle (TCA), and trehalose synthesis are prevalent features of cachexia (Liu *et al*, 2025). Analyzing the expression of the three groups of genes representative of each metabolic pathway, we found a similar sex-specific profile (Fig 6A-C). Among the 24 genes involved in glycolysis, the majority (18 genes) are downregulated in cachectic males, whereas they are unchanged or upregulated in cachexic females (Fig 6B). A comparable trend is observed for genes of the Krebs cycle or trehalose synthesis. Interestingly, we also identified genes exhibiting a reciprocal pattern, being up-regulated or unchanged in females, and downregulated in males (Fig 6A-D). We further investigated the behavior of genes belonging to these three metabolic pathways, using PCA and assessing log2 changes in expression levels between sexes. While PCA did not reveal any specific patterns for trehalose metabolism (Fig 6E), the first principal component of the PCA clearly separated males and females for genes of glycolysis and Krebs cycle (Fig 6F, G). In both cases, cachectic males stand out from healthy males by moving toward the females. (Fig 6F, G). Comparison of correlation coefficients (Fig 6H) also shows that Yki^act^ males are far more similar to Yki^act^ females (r=0,89) than their healthy counterparts (r=0.54), supporting the notion that cachexia strongly influences the metabolism of males harboring intestinal tumors. We further assessed the relative difference in gene expression levels between males and females for each condition (healthy and cachexia, Fig 6I, J). For glycolysis, trehalose and TCA cycle, the sex-specific difference in gene expression is reduced in cachectic conditions, as well illustrated by deregulation of *Pepck1* (Fig 6 D). We therefore conclude that cachexia forces the convergence of expression levels for genes of major metabolic pathways, thereby flattening sex-based disparities.

**Figure 6:**
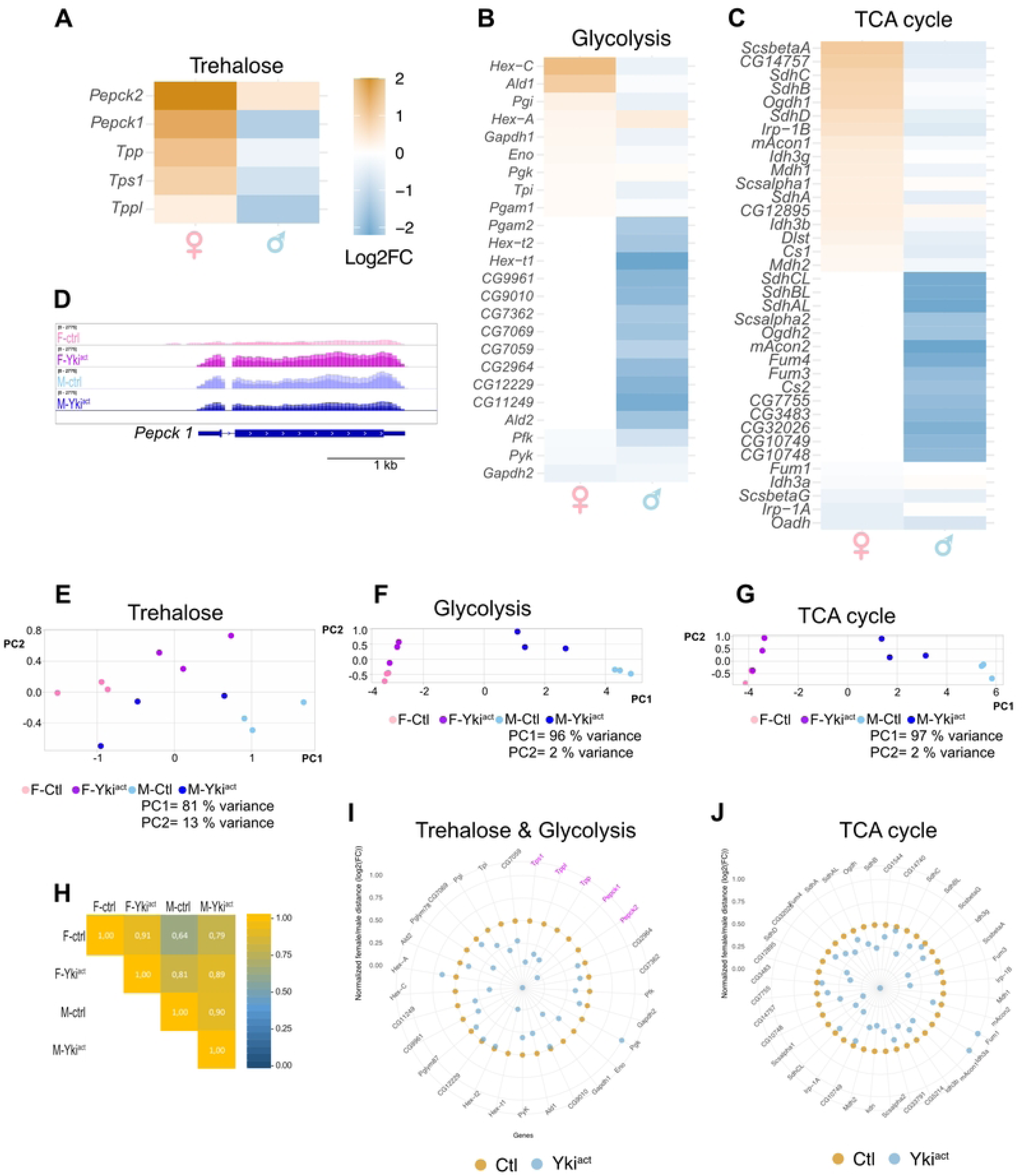
Effect of cachexic state on metabolic pathways. **(A-C).** Heatmaps showing expression dynamics (log2(FC)) of genes involved in trehalose (**A**), glycolysis (**B**), and TCA cycle (**C**), in control and cachexic conditions, for males and females. **(D)**. Genome browser screenshot of the *Pepck1* locus, a representative example of the behavior of genes responsible for the Trehalose metabolism. **(E-G).** PCA of genes involved in the indicated metabolic pathways, in control and cachexic conditions, for males and females. **(H).** Correlation matrixes of gene expression by sex and oncogenic conditions for all metabolic genes. The correlation coefficient is shown in each cell. **(I, J).** Coxcomb plots comparing the relative difference in gene expression between males and females for metabolic genes under control conditions and in response to Yki^act^. (**I**) shows genes involved in glycolysis and trehalose metabolism (pink), and (**J**) genes involved in TCA (Krebs) cycle. The difference was normalized (100%) for the absolute value observed under normal conditions.

## DISCUSSION

One of the main challenges in developing personalized treatments of cachexia is identifying specific tumor characteristics and take sex differences into account. Using a well-established model of intestinal tumor-induced cachexia in *Drosophila* (Liu *et al*, 2022), we investigated here the sex differences in the onset of this syndrome. Using a transcriptomic approach of tumoral intestines and wasting organs included in the body-proper we show that cachexia develops in both sexes, despite initial sex-difference in intestinal stem cell turnover (Hudry *et al*, 2016b). We find that, in both sexes, cachexia is characterized by a state of inflammation within the body and affects the same processes. Although its physiological and transcriptional manifestations differ by sex, the cachectic syndrome blurs initial sex-specific differences and leads tumorous animals toward a comparable cachectic state.

While male and female fly intestines share fundamental functions such as nutrient intake and pathogen defense, they also display distinctive features visible at the tissue and cellular levels (Blackie *et al*, 2024). After mating, the female’s intestine undergoes significant changes in growth, metabolism, and digestive function to meet the increased nutritional demands associated with reproductive function (Kandelouei *et al*, 2026). Consequently, mated-female intestines are longer than those of males and metabolic traits exhibit prominent sex-specific differences. For example, in males, citrate metabolism is influenced by signals from the gonads, impacting food intake and sperm production (Hudry *et al*, 2019). In mated females, the steroid hormone ecdysone, synthesized by the ovaries, stimulates the proliferation of stem cells to support the increased intestinal cell turnover (Ahmed *et al*, 2020; Al Hayek *et al*, 2021). These intrinsic and extrinsic differences profoundly influence the intestinal response to oncogenic stress. Indeed, females are more prone to developing intestinal tumors than males (Hudry *et al*, 2016b), which explain why cachexia following intestinal tumors has primarily been studied in females (Kwon *et al*, 2015). Here, we show that expression of the active form of Yki in ISCs, is sufficient to induce intestinal tumors in both sexes. A notable characteristic of Yki^act^, compared to other oncogenes such as RasV12, Ovo, or Notch, is that it also induces stem cell heterogeneity, rather than merely promoting their hyperproliferation (Pranoto *et al*, 2023). These hybrid ISC/EC cells, probably polyploid, could enhance the secretion of tumor-derived factors, such as Impl2, explaining why male ISCs respond to Yki^act^ but not, or only weakly, to other oncogenes. Furthermore, the heterogeneity of tumor cells is likely to alter the microenvironment and could promote the oncogenic response of male ISCs. In the future, the use of intestinal tumors in males could help elucidate the impact of heterogeneity on tumor development.

In vertebrates, the etiology of cachexia is closely linked to an inflammatory state (Ferrer *et al*, 2023). To date, the production of the cytokine Upd3, an ortholog of IL6 in mammals, has been associated with the development of cachexia in female flies (Ding *et al*, 2021). Our results reveal that Upd3, as Pvf1, ITP and Impl2, are also produced by the male tumorous intestine. It appears that an inflammatory state, which is consistently associated with the development of cachexia in various organisms and models, also develops in *Drosophila*. We observed a robust inflammatory response in both sexes, originating from the wasting organs, most likely from the fat body that is capable of producing AMPs (Lemaitre & Hoffmann, 2007). Several AMPs have been detected in the hemolymph of females cachectic flies, corroborating our findings (Singh *et al*, 2025). The study of Singh and colleagues presents evidence supporting that AMPs are first produced in the intestine in response to epithelium damages due to tumor development. Although our analysis didn’t detect the production of AMPs in the intestine, it remains possible that AMP deregulation might transiently occur in tumors, , and is then relayed, sustained, and amplified by adipose tissues.

In both humans and flies, cachexia profoundly affects the metabolism of distant organs, leading to the wasting of muscle and adipose tissues. Our data confirm that cachexia alters the behavior of these organs in male and female flies, but with some differences in the nature and intensity of the response. For instance, renal failure manifested as abdominal bloating and damages to Malpighian tubules is less severe and/or occurs later in males. Similarly, muscle activity measured by climbing tests is more affected in females (Fig. 3C). Sex-based differences in the intensity of cachexia symptoms have also been reported for anorexia, with fly males being this time affected earlier than females (Petsakou *et al*, 2025). At first glance, our data show that cachexia affects both male and female flies, but with disparities between sexes. However, detailed analyses of deregulated genes in atrophied target organs reveal that these apparent disparities fade during cachexia. Gene expression levels in the body of males and females converge toward a similar cachectic state, with sex-specific differences observed under normal conditions diminishing under cachexia. For example, metabolic genes involved in sugar synthesis, exhibit antagonistic deregulation in males and females, yet the discrepancy in gene expression is smaller in cachectic conditions than in controls. We propose that the cachectic state somehow prevails over sex-specific traits.

To our best knowledge, only a few examples of cachexia have been identified to date in adult flies. These include xenografts of oncogenic larval tissue (Figueroa-Clarevega & Bilder, 2015) and tumors of the male reproductive system (Bhattacharya *et al*, 2024), both of which cause organ atrophy in host flies. As in the case of Yki^act^ intestinal tumors, Impl2 is produced by these other tumors and contributes to trigger the cachectic syndrome. In addition to cancer, cachexia can also arise from kidney failures in humans (Hanna *et al*, 2020). Signs of cachexia have been observed in association with kidney lesions in flies, *e.g.*, upon the depletion of stellate cells, a renal cell type playing key roles in osmotic regulation (Denholm *et al*, 2013). It would be interesting to explore whether the common cachectic signature identified here for intestinal tumors in both sexes can occur in different genetic contexts.

In sum, our findings demonstrate that inflammation is a general characteristic of cachexia, which can develop in both males and females by affecting the same processes (muscular, adipose, and metabolic) but in different ways. This is true even for general pathways, such as glycolysis, that can be affected in opposite ways by cachexia, with key genes being deregulated in an antagonistic manner depending on sex. These results therefore suggest that future studies on the human cachexic syndrome, particularly in its early stages, should benefit from gender-specific approaches.

## Material and Methods

### *Drosophila* stocks and husbandry

Flies were cultured using standard cornmeal food (per liter: 17 g inactivated yeast powder, 80 g corn flour, 9 g agar, 45 g white sugar, and 17 ml of Moldex). Fly lines used in this study are: *esg^ts^ (Jiang et al., 2009); UAS-Yki^3SA^* (Oh and Irvine, 2009) Bloomington Drosophila Stock Center (BDSC) #28817, *white^1118^* (#3605). *esg^ts^* virgin females were crossed with the following males: control (*w^1118^*), *UAS-Yki^3SA^* (referred here to as Yki^act^). Conditional expression in ISCs (driven by *esg-Gal4)* was achieved by maintaining *tub-Gal80^ts^* (*esg^ts^*) expressing flies at 18 °C, until adulthood. Eclosed males and females were aged for 3-days, then shifted to 29°C to induce Yki^act^ expression and kept at 29°C for the indicated period of time.

### Immunohistochemistry

Tissues were dissected in 1XPBS, fixed in 4% formaldehyde in PBS for 1 h at room temperature (or overnight at 4°C), then washed for 5 min in PBS containing 0.1% Triton X100 (PBT). Following a 5 min wash in PBT, tissues were blocked in PBT containing 1% BSA for 30 min. Primary antibodies were incubated overnight at 4 °C, using the following dilution: anti-GFP (Acris Antibodies Cat#TP401) 1:200, anti-phospho-Histone H3 (Upstate Cat#TAKMA312B) 1:2000, anti-Disc-large (DSHB Cat#4F3) 1:100, anti-Mef2 (a gift from Eileen Furlong, Heidelberg, Germany) 1:200. AlexaFluor-488, -555 or -641 secondary antibodies (1:500, Thermo Fischer Scientific) and Alexa-conjugated Phalloidin (1:200, Thermo Fisher Scientific) were incubated for 2 h at room temperature. After three washes, tissues were mounted in Vectashield (Vector Laboratories). Adult thorax muscles were prepared as described by (Leroux *et al*, 2023). Pictures were acquired using Leica SPE and SP5 confocal microscopes. Images were processed with Image J and Affinity software.

### RNA sequencing

Ten male and female individuals were dissected ten days after induction and subjected to TRIzol (Ambion) extraction, according to the manufacturer’s protocol. After removing the head, intestines were dissociated from the rest of the body, referred here to as “body proper”, comprising muscles, fat body, Malpighian tubules, sexual apparatus, and other residual tissues. Three independent samples were run for each condition. RNA quality was checked using an Agilent chip (O’Brien *et al*, 2011). RNA samples were sequenced using Illumina NovaSeq 6000 S2 (paired-end, 100 bp long reads, 40 M reads per sample) at Integragen (https://integragen.com). RNA-seq data were recorded, and assigned the reference number GSM9300833-GSM9300856, GSE310415.

### RNA-seq analysis

The quality of each raw sequencing file (fastq) was verified with FastQC v0.12.1 (www.bioinformatics.babraham.ac.uk/projects/fastqc/). Fastq were aligned to the Dm6 version of *D. melanogaster* genome and counted on the annotation from FlyBase database (gtf dmel-all-r6.44) using STAR aligner (STAR_2.5.2b), as well as a genome index constructed using a FASTA file containing coding and non-coding transcripts. Count tables were generated using Salmon v1.9.0 with the keepDuplicates option enabled; all other parameters were kept at default values. Two types of differential expression analyses were performed using the DESeq2 package (v1.46.0). In both cases, DESeq2 intrinsic normalization was applied using the median-of-ratios method.

The first type of analyses consisted of independent comparisons performed separately for males and for females, comparing controls versus Yki^act^ conditions (design = ∼condition). Genes with no or negligible expression were filtered off, retaining those with a sum of normalized counts ≥ 1000 across all samples, within a given sex and tissue. Genes were considered differentially expressed if log2 fold change (log2FC) was > 0.5 or < -0.5, with an adjusted p-value (padj) < 0.05.The second type of analyses corresponded to a global model integrating both male and female datasets. This approach allowed simultaneous evaluation of the effect of the condition (control *vs* Yki^act^) and the differences in deregulation between sexes (males *vs* females), using the following design: design = ∼ sex + condition + sex:condition. Lowly expressed genes were filtered off using the same criterion as above.

This model was not used to identify sex-specific DEGs, but rather to detect genes showing a significant difference in levels of deregulation between males and females. Results were obtained using results (dds, name = “sexM.conditionYki”), and only genes with an adjusted p-value (padj) < 0.05 were considered significant. The reference levels for this analysis, was set to female controls.

Gene categories of interest were defined by comparing gene lists obtained from differential expression analyses (*i.e.*, genes identified as deregulated in males in the independent analysis, genes deregulated in females in the independent analysis, and genes showing a significant difference in deregulation between sexes in the global model).

The first step consisted of comparing male and female gene lists from the independent analyses to distinguish:

i. genes deregulated only in one sex (male-specific or female-specific)
ii. genes deregulated in both sexes.

The gene list obtained from the global model (here referred to as “in interaction”) was used to further refine these categories by identifying the genes whose levels of deregulation significantly differed between males and females.

This approach defined four distinct groups:

- Group MDO: Genes identified as DE in males (independent analysis), with significantly different deregulation levels when compared to females (according to the interaction model).
- Group FDO Genes identified as DE in females (independent analysis) with a significantly different deregulation levels compared to males (according to the interaction model).
- Group CSD: Genes identified as DE in both sexes (independent analyses), and showing significantly different deregulation levels between males and females (according to the interaction model).
- Group TCD: Genes identified as DE in both sexes, with no significant difference in the levels of deregulation between males and females (corresponding to genes deregulated in both sexes with a similar level of deregulation).

### Biostatistics statistical analyses

#### Analysis of Cell Types and Organs

Marker gene lists specific to organs (body proper analysis) and cell types (intestine analysis) were obtained from *Drosophila* single-cell atlases (Cell Atlas (Li *et al*, 2022) and Midgut Cell Atlas (Hung *et al*, 2020)). Marker tables were imported, and only genes associated with a single organ or cell type were retained.

Raw counts for these markers obtained from DESeq2 were transformed using the Variance Stabilizing Transformation (*VST; blind = TRUE*), and then centered and scaled (z-score). For each sample, a marker expression score was calculated for each organ or cell type as the mean of z-scores across all its specific marker genes.

Scores were visualized using boxplots with individual sample points overlaid, allowing comparison of organ- or cell-type–specific expression patterns across experimental conditions. When multiple cell types or organ subtypes could be grouped into a broader category (*e.g.*, pEC, mEC, dEC grouped as EC), they were combined for visualization and interpretation.

#### Scatter plot, histogram & cachexia markers boxplot

The data used to generate the scatter plots, heatmaps, were transformed using the Variance Stabilizing Transformation (*VST; DESeq2, with* blind *= TRUE*) in order to stabilize the variance of count data prior to visualization.

#### Cell type or organ markers enrichment (gsea)

Differentially expressed (DE) genes were ranked by decreasing log2(FC) and used for enrichment analysis based on adult midgut cell types and immune response (GO:0006955). Marker gene lists obtained from single-cell atlases were cleaned to remove missing entries. Multilevel GSEA was performed using the fgsea package (v1.32.0) with 10,000 permutations and *eps* = 0 to compute normalized enrichment scores (NES) and adjusted p-values. Results were visualized as enrichment curves displaying ES, NES, and padj for each cell type.

#### Gene Ontology

DE genes were analyzed for enrichment in Gene Ontology Biological Processes (GO-BP) using clusterProfiler (OrgDb = *org.Dm.eg.db*, v3.20.0; FlyBase gene identifiers; Benjamini–Hochberg correction; *padj* < 0.05, *q* < 0.05; background = all tested genes). Redundant GO terms were removed. The analysis was performed separately on the full DE gene set.

In a same way, DEGs were analyzed by GO Biological Process (GO-BP) GSEA using clusterProfiler (*gseGO; OrgDb = org.Dm.eg.db*; FlyBase identifiers; fgsea method with 10,000 permutations, *eps = 0*). Redundant GO terms were filtered off using simplify (*cutoff = 0.6*).

#### Metabolism analyses

Gene lists corresponding to the selected metabolic categories were retrieved from FlyBase. For each gene, log2(FC) values from male-vs-female differential expression analyses were extracted separately for control and cachectic conditions. To facilitate comparison, male-*vs*-female log2(FC) in control conditions was normalized to 100% for each gene, and corresponding log2(FC) in cachectic conditions was reported as the ratio. Normalized values were visualized as circular histograms allowing comparison of the relative magnitude of sex-specific differences in expression levels between control and cachectic states.

#### Software

FIJI and Affinity packages were used for image analysis and figure construction. R and R studio were used to treat, analyze and generate graphs from RNAseq data. *We used Mistral Medium 3.5 (Mistral AI, 2024) to proofread parts of the text. After using Mistral AI, the authors reviewed and edited the content as needed and take full responsibility for the content of the publication*.

## Data and code availability

RNA-seq, have been deposited at GEO and are publicly available as of the date of publication. Accession numbers are listed in series GSE310415 and avalable in the key resources table.

*To review GSE310415, go to* https://www.ncbi.nlm.nih.gov/geo/query/acc.cgi?acc=GSE310415 *Enter token absvwgysbnybnyr into the box*.

## Code availability

This paper does not report original code. Scripts used on this paper are available on https://src.koda.cnrs.fr/cachexia/sex-disparities

Any additional information required to reanalyze the data reported in this paper is available from the lead contact upon request.

## Acknowledgements

We thank Marion Aguirrebengoa (BigA platform, CBI) for her bioinformatic help and advices. We thank Michèle Crozatier for discussions and critical reading of the manuscript. This work was supported by grants from the Agence Nationale de la Recherche to J. Z. (ANR-17-CE13-0002), Canceropole GSO-emergence N°2023-E10 and Fonroga Fundation. A.R. was supported by fellowships from Ligue contre le Cancer.

## Author contributions

Conceptualization: Jennifer Zanet, Cédric Polesello

Data curation: Alexia Rivero, Cédric Polesello.

Funding Acquisition: Jennifer Zanet

Formal analysis: Alexia Rivero.

Investigation: Alexia Rivero, Margaux Techer, Nourhene Ammar, François Payre, Hadi Boukhatmi, Jennifer Zanet and Cédric Polesello.

Methodology: Alexia Rivero

Project Administration: Hadi Boukhatmi, Jennifer Zanet and Cédric Polesello

Resources: Hadi Boukhatmi, Jennifer Zanet and Cédric Polesello

Software: Alexia Rivero

Supervision: Hadi Boukhatmi, Jennifer Zanet and Cédric Polesello

Visualization: Alexia Rivero, Margaux Techer, Nourhene Ammar, François Payre, Cédric Polesello Writing – original draft: Cédric Polesello

Writing-Review & Editing: Alexia Rivero, Nourhene Ammar, François Payre, Hadi Boukhatmi, Jennifer Zanet, Cédric Polesello

